# Identifying long-range synaptic inputs using genetically encoded labels and volume electron microscopy

**DOI:** 10.1101/2021.12.13.472405

**Authors:** Irene P. Ayuso-Jimeno, Paolo Ronchi, Tianzi Wang, Catherine Gallori, Cornelius T. Gross

**Affiliations:** Epigenetics & Neurobiology Unit, European Molecular Biology Laboratory (EMBL), Via Ramarini 32, 00015 Monterotondo (RM), Italy; Electron Microscopy Core Facility (EMCF), European Molecular Biology Laboratory (EMBL), Meyerhofstr 1, 69117, Germany

## Abstract

Enzymes that facilitate the local deposition of electron dense reaction products have been widely used as labels in electron microscopy (EM). Peroxidases, in particular, can efficiently metabolize 3,3’-diaminobenzidine tetrahydrochloride hydrate (DAB) to produce precipitates with high contrast under EM following heavy metal staining, and can be genetically encoded to facilitate the labeling of specific cell-types or organelles. Nevertheless, the peroxidase/DAB method has so far not been reported to work in combination with 3D volume EM techniques (e.g. Serial blockface electron microscopy, SBEM; Focused ion beam electron microscopy, FIBSEM) because the surfactant treatment needed for efficient reagent penetration disrupts tissue ultrastructure and because these methods require the deposition of large amounts of heavy metals that can obscure DAB precipitates. However, a recently described peroxidase with enhanced enzymatic activity (dAPEX2) appears to successfully deposit EM-visible DAB products in thick tissue without surfactant treatment. Here we demonstrate that multiplexed dAPEX2/DAB tagging is compatible with both FIBSEM and SBEM volume EM approaches and use them to map long-range genetically identified synaptic inputs from the anterior cingulate cortex to the periaqueductal gray in the mouse brain.

## Introduction

A major challenge in circuit neuroscience remains the reliable determination of neuron-to-neuron connectivity in the brain of laboratory animals. Long-range neuronal projections can be visualized using anterograde transported fluorescent labels or genetically-encoded fluorescent proteins. However, the inability to resolve synaptic structures by diffraction-limited light microscopy methods makes it challenging to identify the specific target cells that receive such long-range connections, and electron microscopy remains the method of choice to reliably identify synaptic contacts. In particular, the anterograde axonal transport of horseradish peroxidase (HRP) has been widely used in conjunction with the substrate DAB to label axonal processes, identify synaptic targets, and infer their neurotransmitter identity (i.e. symmetric vs. asymmetric)^1–5^. In more recent variants of this method, HRP has been delivered by viral vectors in a cell-type specific manner, allowing for the mapping of specific genetically-defined axonal projections. In one such approach, HRP was fused to the neurotransmitter vesicle protein synaptobrevin to selectively label axonal boutons^6^ and similar fusion tags have been developed based on the miniSOG tag^7,8^.

However, at least two methodological barriers limit the identification of synaptic contacts using existing genetically-encoded EM tagging methods. First, these labeling methods do not allow for multiplexed tagging and thus cannot be used to map synaptic connectivity between genetically defined pre- and postsynaptic neurons. Second, these labeling methods are incompatible with volume EM approaches required for the comprehensive and high-throughput assessment of synaptic architectures in tissues because of the poor tissue penetration of the DAB substrate^9^. Fortunately, both of these impediments appear to have been resolved by the recent development of a peroxidase variant, dAPEX2, with enhanced enzymatic activity^10^. dAPEX2 is derived from soybean ascorbate peroxidase and was shown to successfully deposit EM-visible DAB reaction products in tissue sections at up to 200 microns depth as visualized after sectioning by transmission EM^10^. These data suggest that dAPEX2 may be compatible with blockface volume EM methods that require homogeneous DAB reagent infiltration into thick tissue samples prior to sectioning.

Here we demonstrated the multiplex imaging of synaptic contacts between long-range excitatory inputs and genetically-defined target neurons using dAPEX2 and volume EM in laboratory mice. We showed that mitochondrial matrix and endoplasmic reticulum-tagged dAPEX2 can be reliably detected in thick tissue samples by blockface EM methods and used to systematically identify pre- and postsynaptic contacts in the resulting 3D isotropic imaging datasets. To demonstrate the power of this approach we applied it to uncover the cellular targets of long-range corticofugal projections from the mouse anterior cingulate cortex (ACC) to the dorsal periaqueductal gray (dPAG). We chose ACC-dPAG projections^11–15^ for study because ACC layer 5 excitatory projections have been shown to inhibit neural activity in dPAG and suppress its behavioral output^14,15^ despite functional channelrhodopsin circuit mapping showing that ACC excitatory neuron afferents excite glutamatergic, but not GABAergic neurons in dPAG^11–14^ (13% of dPAG *Vglut2*+ neurons receive monosynaptic excitatory inputs from ACC)^14^. These apparently paradoxical findings could be explained, however, by the observation that virtually all glutamatergic dPAG neurons showed a ChR2-dependent tonic reduction in frequency of spontaneous excitatory postsynaptic currents (sEPSCs), suggesting that ACC inputs act widely to suppress the probability of release of excitatory inputs to *Vglut2*+ dPAG neurons via a presynaptic neuromodulatory mechanism^14^. We hypothesized that such modulation could be mediated either by direct inhibitory axon-axonic contacts of glutamatergic ACC projections onto excitatory boutons/axons that innervate glutamatergic dPAG neurons, or via the feedforward activation of local inhibitory neurons that make inhibitory axon-axonic contacts, and set out to use multiplex dAPEX2/DAB labeling and volume EM to distinguish these possibilities. Our ultrastructure study allowed us to confirm the selective enervation of excitatory neurons in dPAG by ACC, rule out the formation of direct axon-axpnic inputs, and implicate non-glutamatergic/non-GABAergic neuromodulatory neurons in cortical feedforward inhibition.

## Results

### dAPEX2 visualization by volume electron microscopy

To determine whether multiplex pre- and postsynaptic dAPEX2 labeling could be used to visualize genetically defined synaptic contacts by volume EM, we expressed mitochondria-targeted dAPEX2 (Matrix-dAPEX2)^10^ in the mouse ACC and endoplasmic reticulum-targeted dAPEX2 (ER-dAPEX2)^10^ in dPAG. Matrix-dAPEX2 (AAV1/2-*Ef1α*::COX4-dAPEX2) was used for labeling ACC axons because mitochondria are routinely found in axonal varicosities^16^, while ER-dAPEX2 was used for labeling dPAG cell bodies as endoplasmic reticulum is found widely in both soma and dendrites^17^. ER-dAPEX2 expression was restricted to glutamatergic or GABAergic target cells in dPAG using a Cre-dependent adeno associated virus (AAV) expressing EM-dAPEX2 (AAV1/2-*Ef1α*::DIO-IGK-dAPEX2-KDEL) delivered to *Vglut2*::Cre or *Vgat*::Cre mice, respectively (**Fig. 1a**, **Table S1**). We confirmed that the two dAPEX2 tags were effectively expressed in ACC and dPAG using light microscopy. Following DAB staining, frontal cortex brain sections from the infected animals showed a dark variegated precipitate across the injection site in ACC and fiber-like staining in its targets, including nucleus accumbens (NAc) shell, claustrum (CL) and insular cortex (IC)^13,14,18,19^ (**Fig. 1b**, left). In dPAG a pattern of staining corresponding to that described for ACC projections^1^ was observed caudally to the injection site (**Fig. 1b**, right). At the center of the infection site in dPAG a pattern of staining corresponded to the localization of ER in cell bodies^17^ was observed (**Fig. 1c**). Next, we confirmed the subcellular localization of dAPEX2 staining in dPAG by transmission EM. Both Matrix-dAPEX2 and ER-dAPEX2 staining were confirmed to be localized to their respective organelles, as reported previously^10^ (**Fig. 1d**). Finally, we tested whether the dAPEX2 label could be visualized by volume EM. Imaging samples by SBEM revealed robust Matrix-dAPEX2 and ER-dAPEX2 labeling (**Fig. 1e**) demonstrating that dAPEX2 labeling is compatible with volume EM.

**Figure 1.**
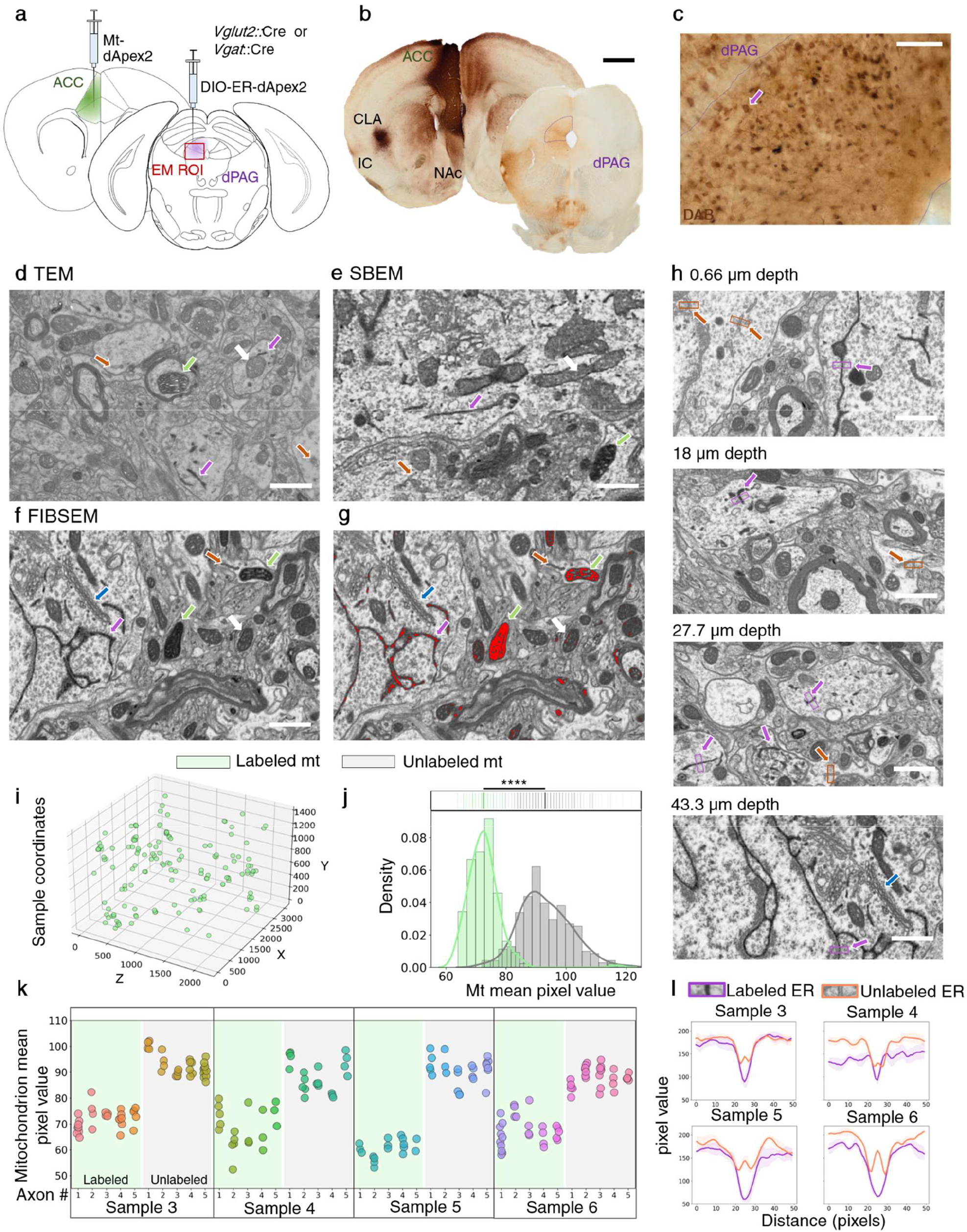
dAPEX2 labels are visible in brightfield, TEM, FIBSEM, and SBEM images. **a** *Vglut2*::Cre or *Vgat*::Cre mice were injected with AAV1/2-*Ef1α*::COX4-dAPEX2 (Mt-dApex2) in ACC and with AAV1/2-*Ef1α*::DIO-IGK-dAPEX2-KDEL (ER-dAPEX2) in dPAG. **b** Representative brightfield images of sections containing ACC and PAG showed a dark precipitate in ACC and projection sites in claustrum (CL), insular cortex (IC), nucleus accumbens (NAc), and dPAG (n = 6 animals; scale bar is 1mm). **c** Detail of PAG infection site. Somas appear DAB stained. Purple arrow points to putative ER-dAPEX2 infected cell (n = 6 animals, scale bar is 100 μm). **d** ER and mitochondria directed dAPEX2 is visible in a brain sample acquired with TEM (n = 2 samples from one animal; Scale bar is 1 μm for **d-h**). **e** ER and mitochondria directed dApex2 is visible in a brain sample acquired with SBEM (n = 1 animal). **f** ER and mitochondria directed dAPEX2 is visible in a brain sample acquired with FIBSEM (n = 4 animals). **g** (same sample as **f**) ER and mitochondria directed dAPEX2 is highlighted with a binary mask by pixel thresholding. **h** Examples of ER-dAPEX2 staining at 0.66, 18, 27.7 and 43.3 μm from the surface of the sample. **i** 3D distribution of dAPEX2 labeled mitochondria in a representative FIBSEM sample (sample 3). **j** Kernel density estimation and histogram of mitochondria mean pixel value (n = 148, unlabeled, green; n = 255, unlabeled, gray). **k** Mitochondria mean pixel value in both labeled and unlabeled axons is similar across all mitochondria in one axon (n = 5 axons per sample). **l** Pixel value line plots of labeled vs. unlabeled ER show unimodal or bimodal profiles, respectively across samples (top, n = 5 ER per sample; green arrows: labeled mitochondrion; white arrow: unlabeled mitochondrion; purple arrow: labeled ER; orange arrow: unlabeled ER; blue arrow: golgi apparatus).

To characterize ACC-PAG synaptic contacts in detail we processed samples for FIBSEM, a volume EM method that benefits from isotropic resolution and is well-adapted for the 3D reconstruction of synaptic architecture^20–24^. Both Matrix-dAPEX2 and ER-dAPEX2 labeling could be readily identified in FIBSEM images (**Fig. 1f**). Non-stained ER presented a clear membrane-bounded lumen^8,10^ making the identification of stained ER straightforward and reliable (**Fig. 1l**). Unlabeled mitochondria presented an electron dense matrix^25^ – particularly in samples prepared with the rOTO^26^ protocol – that provided high contrast and could pose difficulties for distinguishing matrix stained mitochondria. To facilitate the reliable identification of labeled mitochondria we developed a binary mask based on a pixel value threshold set to reference-stained mitochondria. The binary mask was used to classify mitochondria and ER as stained or unstained (**Fig. 1g**). Following selection of mitochondria according to morphological criteria stained mitochondria were identified as those with 1) darker mitochondrial matrix compared to unstained mitochondria, and 2) light cristae similar to unstained mitochondria. Next, we asked whether dAPEX2 staining was homogeneous throughout the thickness of the sample. Both ER-dAPEX2 and Matrix-dAPEX2 labels could be observed throughout the volume of a representative sample (**Fig. 1hi**, **Fig. S1**). To quantify homogeneity across the sample we averaged the pixel values of a representative 2D plane of each mitochondrion. The mean pixel value of all selected stained mitochondria was significantly different from the mean pixel value of unstained mitochondria selected from a representative XY and XZ plane (p-value = 4.76 ×10^−95^; **Fig. 1j**). Moreover, when multiple axons containing labeled and unlabeled mitochondria were examined mitochondria were found to be consistently classified within a single axon (**Fig 1k**; N = 40 axons, 5 labeled and 5 unlabeled axons/sample). On rare occasions, we observed instances of dendrites that contained putative dAPEX2-stained mitochondria (**Fig. S2**). Although the origin of this labeling is not clear, it is unlikely to be due to retrograde transport of AAV in our experiments as dPAG is not known to project to ACC, and may instead be related to the capacity of serotype AAV1 virus particles to transynaptically infect neurons in target tissues, albeit at low efficiency^27^. Finally, no ER or mitochondria labeling was observed in control samples obtained from non-infected animals (**Table S1**). In these samples the quality of labeling and average pixel value of mitochondria was comparable to unstained mitochondria in the stained samples (**Fig. S3**).

### ACC projections target specific neuron sub-types in dPAG

Having shown that the multiplex dAPEX2/DAB method could be used to reliably identify axons and dendrites by volume EM we turned to applying the method to identify the synaptic targets of long-range ACC projections in dPAG. Initially, selected axons containing labeled mitochondria (N = 2) and dendrites containing labeled ER (N = 24) were fully traced in one representative isotropic FIBSEM sample (Sample 3; see **Table 1** and **Fig. S4** for full experimental inventory; **Fig. 2a-c**, **Fig. S1**, **Table S3**). Labeling was confirmed as reliable in this dataset as the mean pixel value of labeled mitochondria differed significantly from that of mitochondria contained in nearby putative unlabeled axons (N = 2, 8, and 4 mitochondria/axon, P-value = 3.55×10^−13^; **Fig. 2d**). The majority of labeled dendrites (N = 17/24) were aspiny (**Fig. 2a**; **Fig. S1**; **Table 1, Fig. S4**), while the remaining labeled dendrites were spiny (N = 1.7±0.81 spines/dendrite). Representative labeled axons (N = 2) made exclusively excitatory synapses (symmetric synapses containing a postsynaptic density, PSD; **Fig. 2a-c**) with no evidence for axon-axonic synapses. Next, a further 46 axons containing labeled mitochondria were traced and analyzed in samples 3, 4, 5 and 6. The analyzed 48 ACC axons contained 67 synaptic boutons establishing contact sites (**Table 1**, **Fig. S4**). These were identified in the sample and classified based on the type of synapse they established (symmetric vs. asymmetric) and whether the postsynaptic dendrite was labeled (**Fig. 3ab**). Analyzed ACC axons contained on average 3.3±2.7 labeled mitochondria (**Table S2**). The presence of sparse mitochondria facilitated the task of reconstructing axons in 3D, as the axon could be traced from multiple starting points at labeled mitochondria. The vast majority of labeled synapses (N = 63/67, 94%) established asymmetric, putative excitatory synapses^28^, while one labeled bouton established a synapse that appeared symmetric (**Fig. 3c**). This result aligns with previous anatomical and electrophysiological data^3^ demonstrating that ACC neurons projecting to dPAG are layer 5 glutamatergic pyramidal neurons. At this point we cannot explain the presence of the putative inhibitory synapse, as retrograde labeling experiments found no evidence for GABAergic neurons in ACC that project to PAG^14^. However, because the remaining synapse in this axon was asymmetric (**Table 1**, contact sites 20.1 and 20.2, **Fig. S4**) we suspect this represented a false negative.

**Figure 2.**
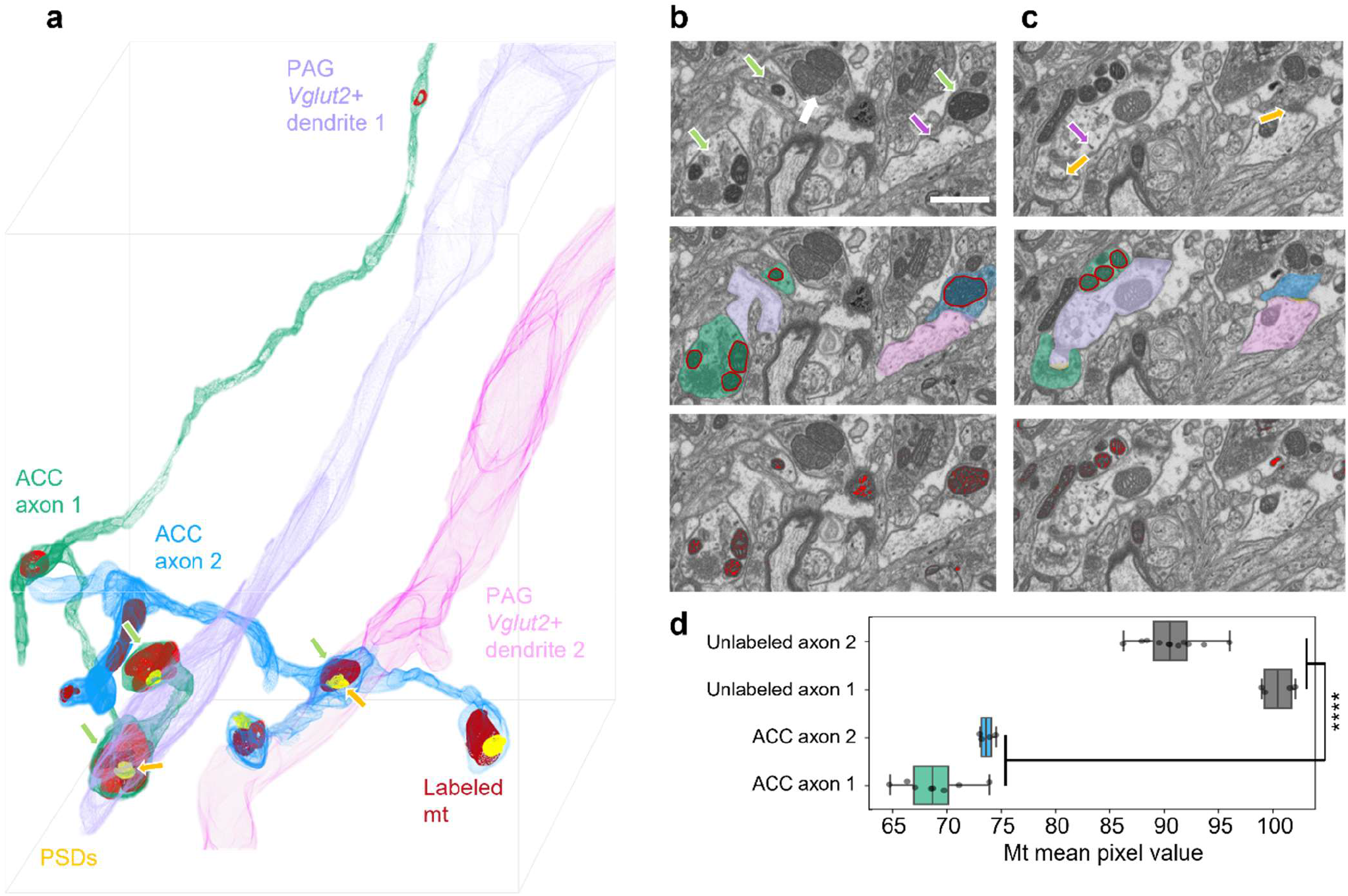
dAPEX2 labeled axons and dendrites can be reconstructed in 3D. **a** 3D reconstruction of two representative ACC axons (blue and green) containing several labeled mitochondria (mt). Both axons established exclusively asymmetric synapses with postsynaptic densities (PSDs). Two representative aspiny *Vglut2*+ target dendrites were reconstructed. **b** Representative planes showing dAPEX2 labeled ACC axons and dendrites. Top: labeled mitochondria and ER are indicated. Middle: segments are reconstructed. Bottom: binary mask highlights labeled mitochondria (scale bar is 1 μm for **b-c**). **c** Same ROI at different depth showing asymmetric synapses with PSDs. **d** Mitochondria mean pixel value for labeled axons was significantly lower than for unlabeled axons (t-test, P = 3.55×10^−13^, sample 3; green arrow: labeled mitochondria; white arrow: unlabeled mitochondria; purple arrow: labeled ER; yellow arrow: PSD).

**Figure 3.**
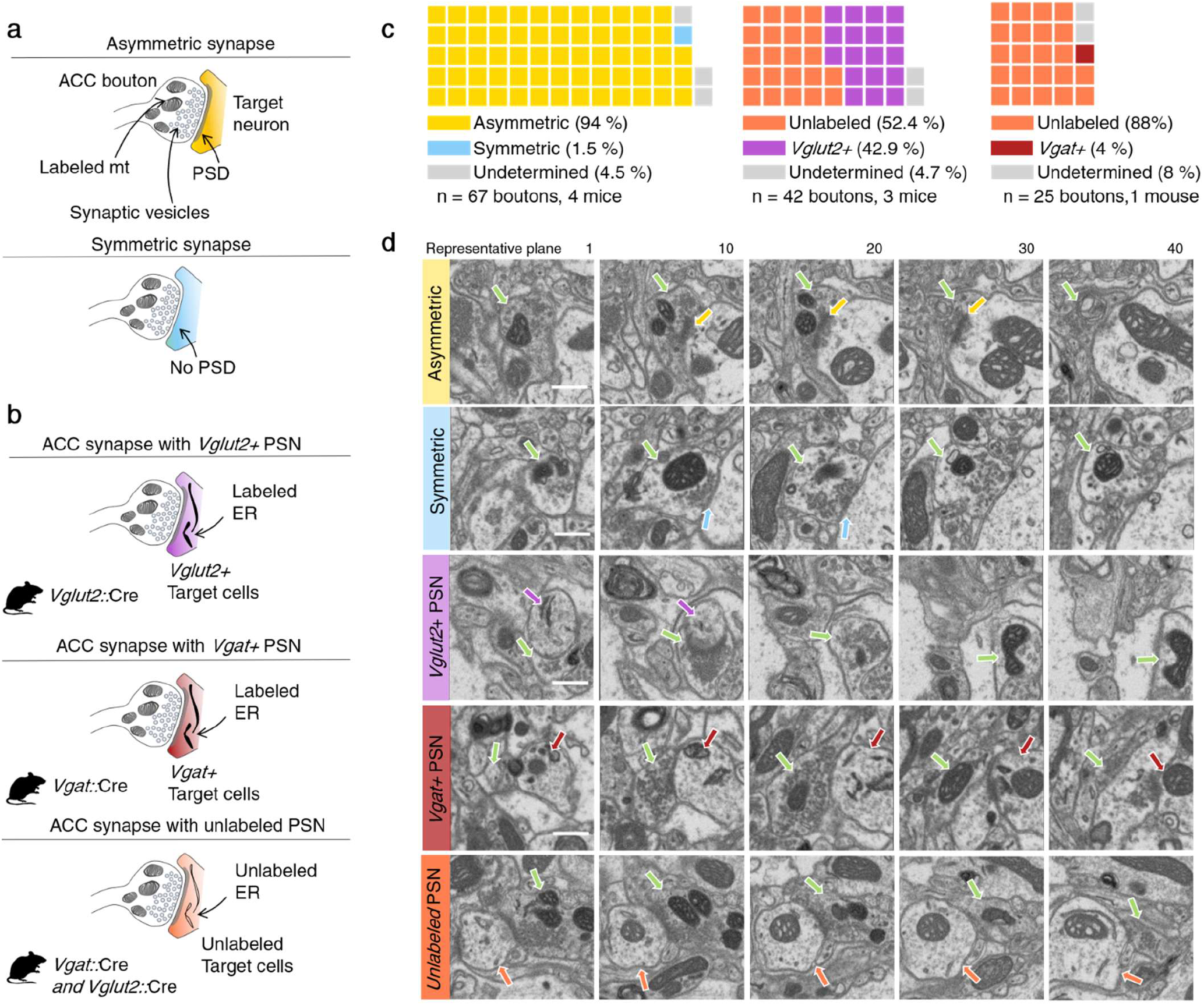
Identification of ACC targets in PAG. **a** Diagram indicating the features of asymmetric (with PSD; putative glutamatergic) and symmetric (without PSD; putative GABAergic) synapses. **b** Diagram indicating the three categories of ACC-dPAG synapses found. **c** Left: Across all mice 94% (63/67) of ACC boutons established asymmetric and 1.5% (1/67) symmetric synapses. Middle: in *Vglut2*::Cre mice 42.9% (18/42) of ACC boutons established synapses with *Vglut2*+ cells, while 52.4% (22/42) of them established synapses with unlabeled cells. Right: 4% (1/25) of ACC boutons established synapses with *Vgat*+ cells, while 88% (22/25) established synapses with unlabeled cells. **d** Exemplary consecutive planes of an asymmetric synapse, a symmetric synapse, and contacts with a *Vglut2*+, *Vgat*+, and unlabeled postsynaptic neuron (PSN; scale bar is 1 μm; green arrow: labeled mitochondrion; white arrow: unlabeled mitochondrion; purple arrow: labeled ER in *Vglut2*+ cell; red arrow: labeled ER in *Vgat*+ cell; orange arrow: unlabeled ER; yellow arrow: PSD).

**Table 1.**
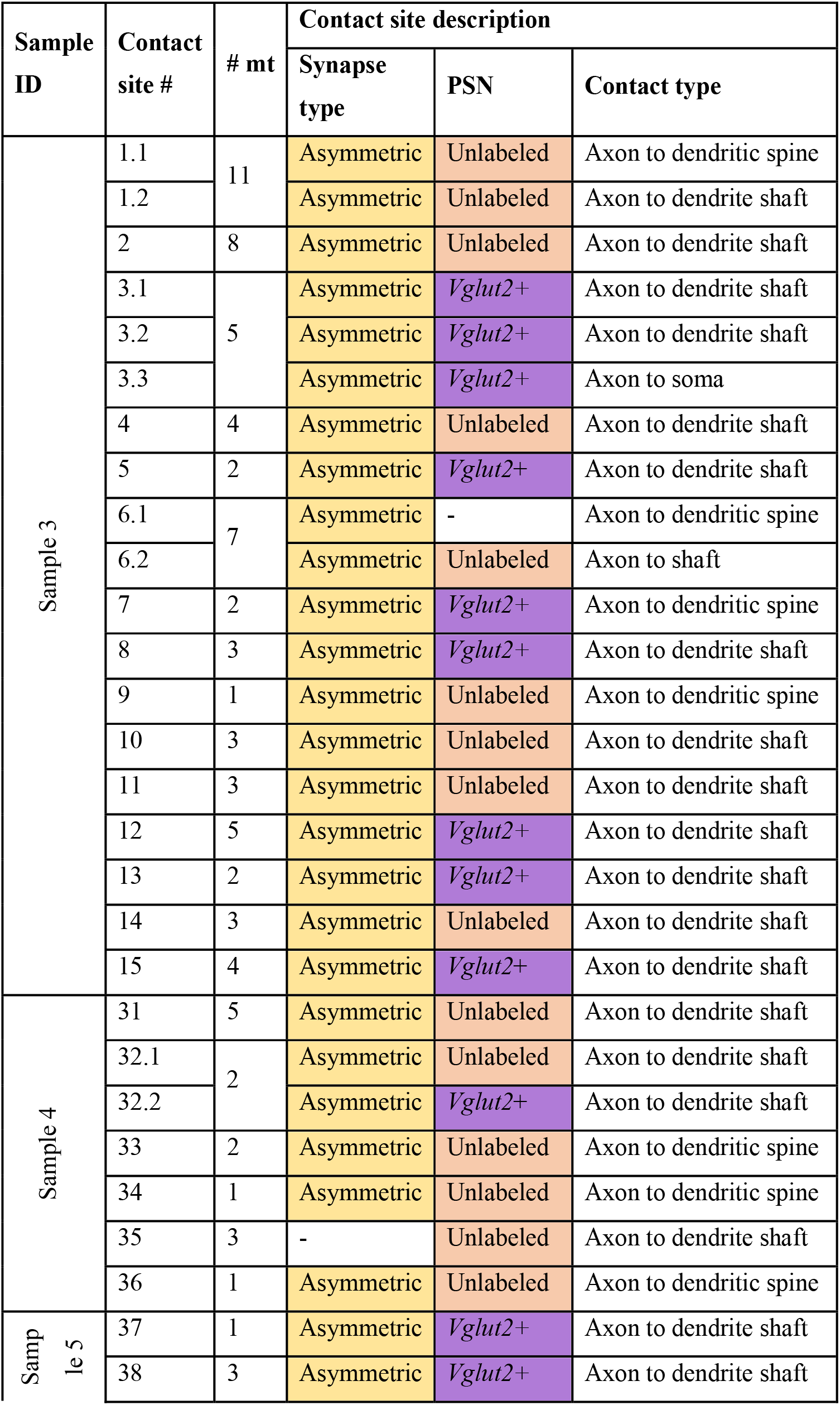
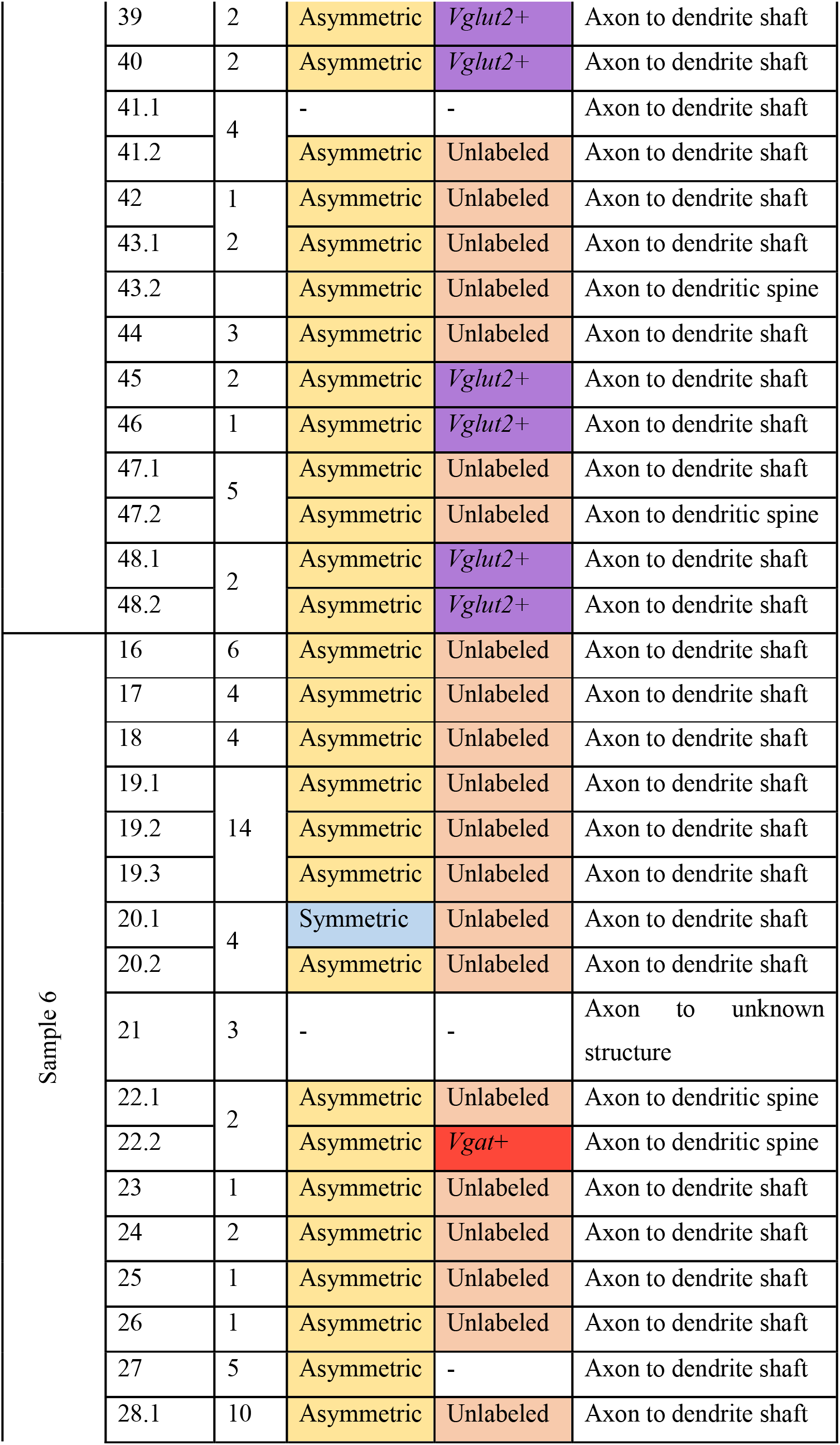
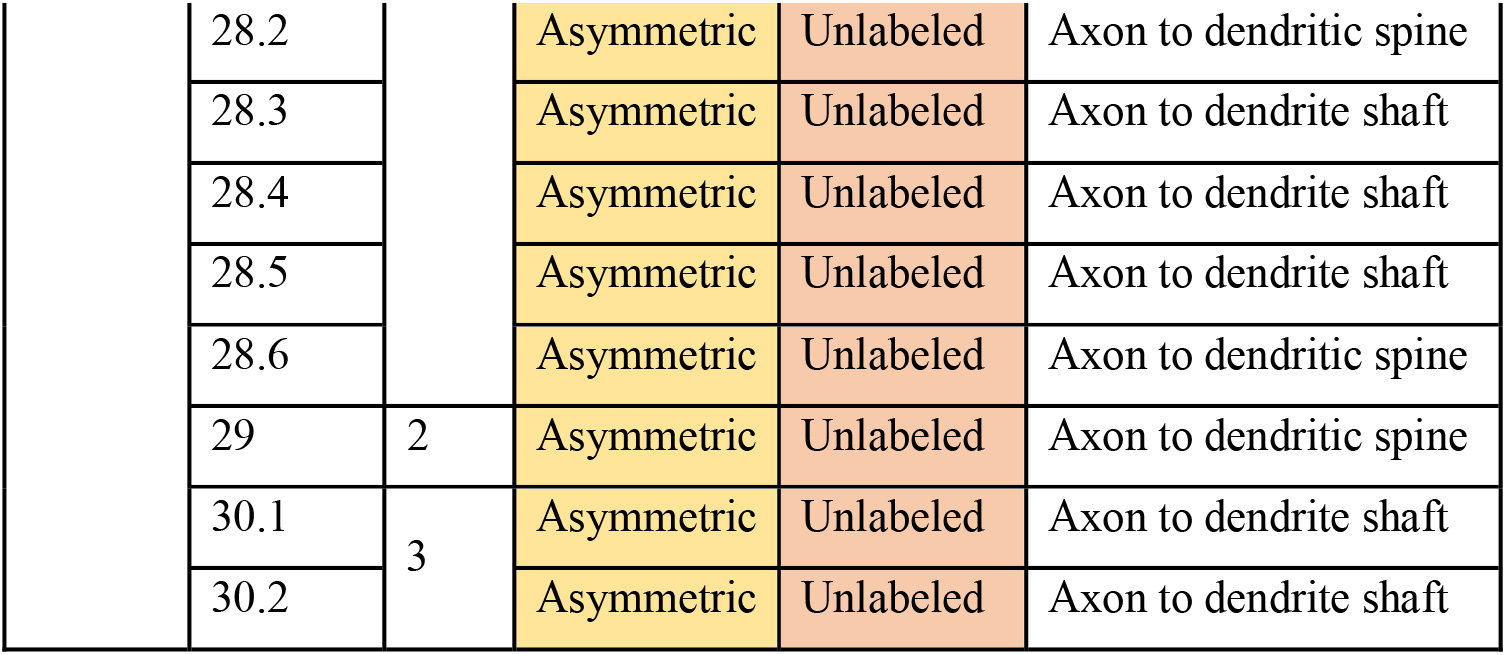
Summary of examined ACC axons in FIBSEM samples (see Fig. S4 for list of samples)

Finally, we explored whether ACC neurons synapsed preferentially onto glutamatergic or GABAergic neurons in dPAG. Constitutive Matrix-dAPEX2 was expressed in ACC and Cre-dependent EM-dAPEX2 was expressed in dPAG of either *Vglut2*::Cre (N = 3) or Vgαt::Cre (N = 1) mice. In *Vglut2*::Cre mice 42.9% (N = 18/42) of putative ACC boutons identified in dPAG synapsed onto *Vglut2*+ neurons and 52.4% (N = 22/42) synapsed onto unlabeled neurons (**Fig. 3 c**). Consistent with data from cell-type specific mono-synaptic rabies tracing^3^ that found no cortical inputs to GABAergic cells in dPAG, in *Vgat*:: Cre mice, 4% (N = 1/25) of putative ACC boutons identified in dPAG synapsed onto *Vgat*+ neurons. These results demonstrate that ACC pyramidal cells do not appreciably target GABAergic neurons, but rather synapse onto a balance of glutamatergic neurons and non-glutamatergic/non-GABAergic, putative neuromodulatory neurons in dPAG.

## Discussion

We have shown that the recently described genetically-encoded EM label dAPEX2 can be effectively combined with volume EM to trace and classify genetically-defined synaptic inputs in mouse brain tissue. Critically, we showed that both mitochondrial and endoplasmic reticulum-trafficked dAPEX2 (Matrix-dAPEX2 and ER-dAPEX2, respectively) could be used to consistently and robustly distinguish pre- and postsynaptic partners across the full depth of tissue blocks routinely used in both SBEM and FIBSEM. To demonstrate the power of combining genetically encoded multiplex dAPEX2 labeling and volume EM we used the method to identify the postsynaptic targets of long-range cortical projections to the brainstem in laboratory mice. Our results show that ACC layer 5 pyramidal neurons make excitatory synapses onto glutamatergic, but not GABAergic neurons in dPAG. Moreover, ACC corticofugal projections make nearly half of their excitatory synapses onto a third, non-glutamatergic/GABAergic population in dPAG. These findings open the possibility that excitatory corticofugal projections may suppress brainstem neuronal activity^14^ via local neuromodulatory feedforward inhibition.

The development of a technology to visualize the full three-dimensional synaptic architecture of pairs of genetically identified neurons in brain tissue will find wide application in circuit neuroscience. Multiplexed volume EM labeling has been possible using correlated light and EM methods (i.e. correlative light and electron microscopy, CLEM), but these are cumbersome and have restricted throughput^29–32^. The advent of light microscopy approaches that reach nanometer resolution, including super-resolution methods^33,34^ and expansion microscopy^35–37^, are likely to bring viable alternatives to EM ultrastructure reconstruction. However, a major advantage of EM over fluorescent labeling of synapses remains its capacity to visualize the full cellular context of labeled structures, although the recent development of unbiased fluorescent staining procedures, such as those that stain the extracellular milieu^38^, could bring similar advantages to light microscopy. Nevertheless, volume EM remains the benchmark method for an unbiased visualization of the ultrastructural environment surrounding synapses and a critical tool for the systematic discovery of brain connectivity.

Our investigation of ACC projections to dPAG provided a test case for the combined application of volume EM and multiplex dAPEX2. Previous work^14^ had established that ACC layer 5 pyramidal cells make long-range excitatory synapses onto glutamatergic, but not GABAergic neurons in dPAG, but nevertheless inhibit neural activity in the target structure by suppressing the frequency of spontaneous excitatory inputs to excitatory dPAG neurons, suggesting that they have a presynaptic neuromodulatory function. Two hypotheses were put forward to explain this inhibitory neuromodulation. Either long-range cortical inputs inhibit presynaptic release probability by making direct axon-axonic synapses onto local excitatory boutons that innervate glutamatergic dPAG targets, or long-range cortical inputs synapse onto local interneurons that in turn inhibit release probability via axon-axonic contacts onto dPAG neurons. Our EM study was able to draw two conclusions. First, we could not find evidence for the formation of axon-axonic contacts by long-range ACC inputs. Of 67 ACC boutons identified, none showed synaptic vesicles in apposition to a bouton that might indicate presynaptic modulation, although it remains possible that we missed such structures, especially if they were rare or apposed to axon terminal segments rather than boutons. *In vitro* electrophysiological studies of the presynaptic neuromodulatory effect showed that it suppresses approximately 50% of the spontaneous excitatory postsynaptic potentials across all glutamatergic neurons recorded^14^, suggesting that the presynaptic modulation must be robust and widespread and making it less likely that a direct axon-axonic neuromodulation would have been missed. Notably, presynaptic ACC neuromodulation was not shown to suppress excitatory inputs to GABAergic neurons, demonstrating the specificity of the phenomenon^14^. Second, we found that ACC inputs consistently targeted both glutamatergic neurons as well as a population of neurons that were neither glutamatergic nor GABAergic (Fig. 3). The identity of this second target population is presently unknown. Neurons that use peptidergic neurotransmission and do not co-release glutamate or GABA have recently been described in the Edinger-Westphal nucleus medial and ventral to the PAG^39^ whose CART+ neurons were shown to modulate fear responses in rodents^39^. Neurons releasing peptidergic neuromodulators have been described in dPAG^40^, including a population that releases enkephalins^41^. Interestingly, enkephalins have been shown to exert presynaptic inhibitory effects on both GABAergic and glutamatergic synapses in PAG, suggesting that they may be mediators of presynaptic feedforward neuromodulatory inhibition^42^. In summary, we have extended the application of a recently developed, enhanced peroxidase-based EM label to volume EM approaches and demonstrated the power of this technique to identify the synaptic targets of long-range corticofugal projections. We expect this method to become a routine part of the circuit neuroscience toolbox.

## Abbreviations

AAV: Adeno associated virus
ACC: Anterior cingulate cortex
CL: Claustrum
CLEM: Correlative light and electron microscopy
DAB: 3,3’-diaminobenzidine tetrahydrochloride hydrate
dPAG: Dorsal periaqueductal grey
EM: Electron microscopy
EMPIAR: Electron Microscopy Public Image Archive
EMBL: European Molecular Biology laboratory
ER: Endoplasmic reticulum
EU: European Union
FIBSEM: Focused ion beam electron microscope
HRP: horseradish peroxidase
IC: Insular cortex
LUT: Lookup tables
Mt: Mitochondria
NAc: Nucleus Accumbens
PAG: Periaqueductal grey
PSD: Postsynaptic density
PSN: Postsynaptic neuron
RT: Room temperature
SBEM: Serial blockface electron microscope
sEPSCs: Spontaneous excitatory postsynaptic currents
TEM: Transmission electron microscope

## Author contributions

I.P.A. and C.T.G. designed research; I.P.A. and P.R. performed research; I.P.A., T.W. and C.G. analyzed data; and I.P.A. and C.T.G. wrote the paper.

## Acknowledgements

We thank the EMBL Rome Laboratory Animal Facility, Genetic & Viral Engineering Facility, Microscopy Facility, and Roberto Voci and Valerio Rossi for support with animal husbandry and management. We thank the EMBL Heidelberg Electron Microscopy Core Facility for supporting volume EM experiments and Dr. Yannick Schwab and Dr. Santiago Rompani for useful comments and advice, Oleksandr Radomskyi for assistance with data analysis, and Martin Schorb for assistance with data processing and sharing. The work was funded by the European Molecular Biology Laboratory (EMBL).

## Materials & Methods

### Animals

All experimental procedures involving the use of animals were carried out in accordance with European Union (EU) Directive 2010/63/EU and under the approval of EMBL Animal Use Committee and Italian Ministry of Health License 541/2015-PR to C.G. Animals were singly housed in temperature and humidity-controlled cages with *ad libitum* access to food and water under a 12 h/12 h light-dark cycle. C57BL/6J mice were obtained from local EMBL colonies. *Vglut2*::ires-Cre and *Vgat*::ires-Cre mice (JAX stock no. 028863 and 028862) were used in heterozygous state. All mice were on a C57BL/6J congenic background.

### Viral production

pAAV-*Ef1α*::DIO-IGK-dAPEX2-KDEL and pAAV-*Ef1α*::COX4-dAPEX2 were purchased from Addgene as a gift from David Ginty (Addgene plasmid #117183; http://n2t.net/addgene:117183; RRID:Addgene_117183; Addgene plasmid # 117176; http://n2t.net/addgene:117176; RRID:Addgene_117176) and packed in chimeric serotype 1/2 AAV vectors by the EMBL Genetic & Viral Engineering Facility. Virus titers were 1.1×10^14^ and 1.3 ×10^14^ vg/ml, respectively.

### Stereotaxic surgeries

Mice were anesthetized with 5% isoflurane and subsequently head fixed in a stereotaxic frame (RWD Life Science) with body temperature maintained at 37 °C. Anesthesia was sustained with 1 to 2% isoflurane and oxygen. The skull was exposed, cleaned with hydrogen peroxide (0.3% in ddH2O) and leveled. Craniotomy was performed with a handheld drill. AAV1/2-*Ef1α*::COX4-dAPEX2 was infused in the ACC (1.11, 1.71 and 1.41 mm anterior to bregma, 0.45 mm left to bregma and 2.0 mm ventral to the skull surface) and AAV1/2-*Ef1α*::DIO-IGK-dAPEX2-KDEL was infused in the dPAG (4.30 posterior to bregma, 1.50 left to bregma and 2.55 mm ventral to the skull surface, at a 20° lateral angle). Injections were unilateral and ~0.2 μl of virus was delivered in each injection site with a pulled glass capillary (intraMARK, 10-20 μm tip diameter, Blaubrand). After virus infusion the skin was sutured and saline solution and Carprofen (5 mg/kg) administered subcutaneously. Mice were allowed 8 weeks for viral expression.

### Perfusion and sectioning

Tissue was prepared following a protocol described previously^10^ with minor modifications. Mice were anesthetized intraperitoneally with 2.5% Avertin (Sigma-Aldrich) and perfused transcardially with warm (~37 °C) Ames medium (MilliporeSigma) with heparin (MilliporeSigma) followed by warm (~37 °C) 2.5% glutaraldehyde (Electron Microscopy Sciences), 2% paraformaldehyde (Electron Microscopy Sciences) in cacodylate buffer (0.15 M sodium cacodylate [Electron Microscopy Sciences] and 0.04%CaCl_2_ [MilliporeSigma]). Ames medium was gassed with Carbogen for animals 7390, 7930, OBO-017266, and 8845. Brains were collected and post-fixed overnight at 4 °C with the same fixative solution. Brains were washed with cacodylate buffer and embedded in 2% low melting point agarose prior to sectioning. ACC and PAG Coronal 100-150 μm sections were cut on a vibratome (Leica Microsystems) in cacodylate buffer.

### DAB staining and bright field microscopy

ACC and PAG sections were washed 2 × 10 min with cacodylate buffer with 50 mM glycine (MilliporeSigma), 1 × 10 min with cacodylate buffer, and then incubated in 1 mL of DAB (MilliporeSigma; 0.3 mg/mL) in cacodylate buffer in the dark for 30 min at RT. Afterwards, 10 ul of 0.3% H2O2 (MilliporeSigma) in cacodylate buffer were added and let react for 1h in the dark at RT. Sections were placed in 3% glutaraldehyde in cacodylate buffer at 4 °C overnight. PAG, DAB stained brain slices were washed1× 10 min in cacodylate buffer, 1×10 min in 50 mM glycine in cacodylate buffer, and 2×10 min in cacodylate buffer. Then they were placed in a slide, freely floating in cacodylate buffer, covered with a coverglass and imaged with a Leica Thunder Imager Cell Culture microscope. 3-4 PAG slices per mouse containing satisfactory putative ER-dAPEX2 labels, or at ca. Bregma AP −4.30 in the case of control brains, were trimmed around PAG and selected for further staining. Remining PAG slices and ACC slices were washed in ddH2O, mounted with mowiol (Calbiochem) and imaged in an Olympus Slideview VS200 microscope.

### Electron microscopy staining

Selected PAG tissue sections were processed following the rOTO protocol^26^ as described previously^10^. Slices were stained in 2% osmium tetroxide (Electron Microscopy Sciences) in cacodylate buffer for 1 h at room temperature, followed by 2.5% potassium ferrocyanide in cacodylate buffer for 1h, and washed 4 × 5 min in ddH2O. Subsequently, sections were incubated in 1% thiocarbohydrazide (Electron Microscopy Sciences) in ddH2O at 40 °C for 15 min and washed 4 × 5 min in ddH2O. Samples were stained in 2% osmium tetroxide in ddH2O for 1 h at room temperature and washed 4 × 5 min in ddH2O. Sections were counterstained overnight with 1% uranyl acetate in 0.05 M sodium maleate (MilliporeSigma; pH 5.15; in samples 6 and 8, uranyl acetate was diluted in ddH2O). Sections were warmed to 50 °C for 2 h in the uranyl acetate solution and then washed 4 × 5 min in ddH2O. Sections were dehydrated in 30%, 50%, 70%, 90% and 3x 100% ethanol, 5 min each, and in propylene oxide (Electron Microscopy Sciences) for 15 min. Once samples were dehydrated, they were infiltrated for 1 h each with 25%, 50%, 75% and 100% Durcupan ACM resin (Electron Microscopy Sciences). Next, samples were placed on a Durcupan block with a flat surface and covered with ACLAR^®^ 33 C Film (Electron Microscopy Science). Samples were let polymerize for 72 h at 60 °C.

### Transmission electron microscopy (TEM)

Sections (70 nm) were cut from polymerized PAG samples using an ultramicrotome (Leica UC7). Sections were imaged without post-staining using a Philips CM120 Biotwin operated with an acceleration voltage of 100kV.

### Focused ion beam electron microscopy (FIBSEM)

Flat-embedded PAG sections were glued to the lateral side of pre-polymerized blocks. Samples were trimmed to expose the region-of-interest (Leica UC7) using a 90° diamond trimming knife (Diatome cryotrim 90). Samples were mounted on a stub using silver epoxy resin (Ted Pella) with the sections perpendicular to the stub surface so that they were parallel to the milling beam. Samples were gold sputter coated (Quorum Q150RS) and FIBSEM imaging performed with a Zeiss Crossbeam 550 using Atlas3D (Fibics & Zeiss) for sample preparation and acquisition. Briefly, the surface above the region-of-interest (typically 50 × 50 μm) was protected with a platinum coat, deposited with 3 nA beam current. Auto-tuning lines were milled on the platinum surface and a carbon coat was deposited on top. Due to the flat embedding and perpendicular mounting of the section a short polishing step was enough to remove the thin layer of empty resin before exposing the embedded tissue. During the stack acquisition the milling was done with 1.5 nA beam current (at 30 kV). For imaging the SEM was operated at 1.5kV/700pA using an ESB detector (collector voltage 1100V). All stacks were acquired with a 10 nm isotropic voxel size. With these settings we acquired volumes of 15 × 30 × 30/55 μm.

### Serial blockface electron microscopy (SBEM)

Flat-embedded PAG sections were mounted on a pin stub using silver conductive epoxy resin (section parallel to stub surface) and trimmed to the right size around the region-of-interest. The sample was then imaged with a Zeiss GeminiSEM 450 equipped with a Gatan 3View system and controlled by the open-source software package SBEMimage^43^. To reduce charging artifacts we used a focal charge compensation device (Zeiss). Images were taken at 1.5kV/300pA and 1.6 μs dwell time with a pixel size of 10 × 10 nm and 40 nm cutting thickness.

### Image preprocessing

The image stacks were registered using the Fiji^44^ plugin “Linear Stack Alignment with SIFT”^45^ (transformation: translation) or AMST workflow^46^ and averaged in the z-axis to reduce noise and image size, producing a final voxel resolution of 10 × 10 × 20 nm. This resolution was sufficient to identify synaptic vesicles and to trace axons in 3D.

Lookup tables (LUT) were inverted and images were saved as a 8-bit tiff stack. Selected 2D planes were denoised using the Fiji plugin Noise2Void^47^ (https://imagej.net/plugins/n2v) for visualization purposes, but analysis was performed on raw data.

### Data analysis and statistics

Aligned, scaled, and LUT inverted tiff stacks were opened using Fiji^44^ and a binary mask was imposed by thresholding pixel values using the threshold Fiji function. Pixel threshold was selected based on one manually seleted reference stained mitochondrion per sample. Mitochondria average pixel value was extracted by selecting one representative 2D plane for each analyzed mitochondrion, using the ROI manager function in Fiji^44^. Processes containing stained mitochondria and ER were traced and categorized manually. 3D reconstruction of selected processes was carried out using 3DMOD (http://bio3d.colorado.edu/imod/). Data visualization and statistics were done using custom Python scripts (Python Software Foundation). No statistical methods were used to predetermine sample sizes. Sample assignment was not randomized. Data collection and analysis were not performed blind to the conditions of the experiments.

### Data availability

All FIBSEM datasets (samples 3 to control 8; **Table S1**) were deposited as tiff files after preprocessing in the Electron Microscopy Public Image Archive (EMPIAR; EMPIAR-10883)^48^ and uploaded to the interactive viewer MoBIE^49^ (https://github.com/mobie/mobie-viewer-fiji) with bookmarks directing the viewer to all figure captions. MoBIE projects can be visualized with the MoBIE Fiji^44^ plugin (https://imagej.net/plugins/mobie) under MoBIE>Open>Open Published MoBIE Project with the name ‘PAG-dAPEX2-FIBSEM’. TEM, SBEM, and raw FIBSEM data is available upon reasonable request.

