## Supplementary figures and tables for "Identifying long-range synaptic inputs using genetically encoded labels and volume electron microscopy"

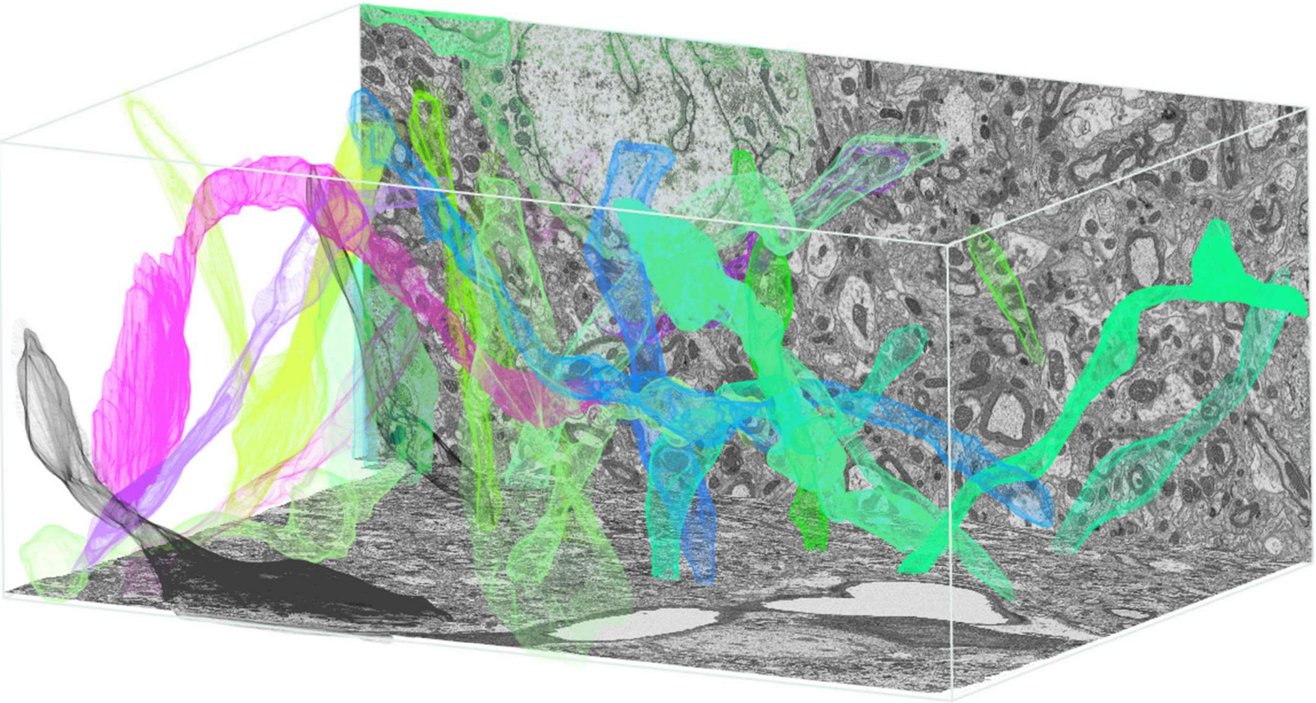

| | With dendritic spines | Without dendritic spines |
|---|---|---|
| With ACC contacts | 1,2,15,18 | 6, 20 |
| Without ACC contacts | 3,11 | 4, 5, 7, 8, 9, 10, 12, 13, 14, 15, 16, 17, 19, 21, 22, 23, 24 |

**Supplementary Figure 1 | ER-stained dendrites are homogeneously distributed in a sample.** Top: 3D reconstruction of 24 ER-dAPEX2 labeled dendrites across the entire sample volume (Sample 3). Bottom: most labeled dendrites were aspiny and did not receive ACC inputs.

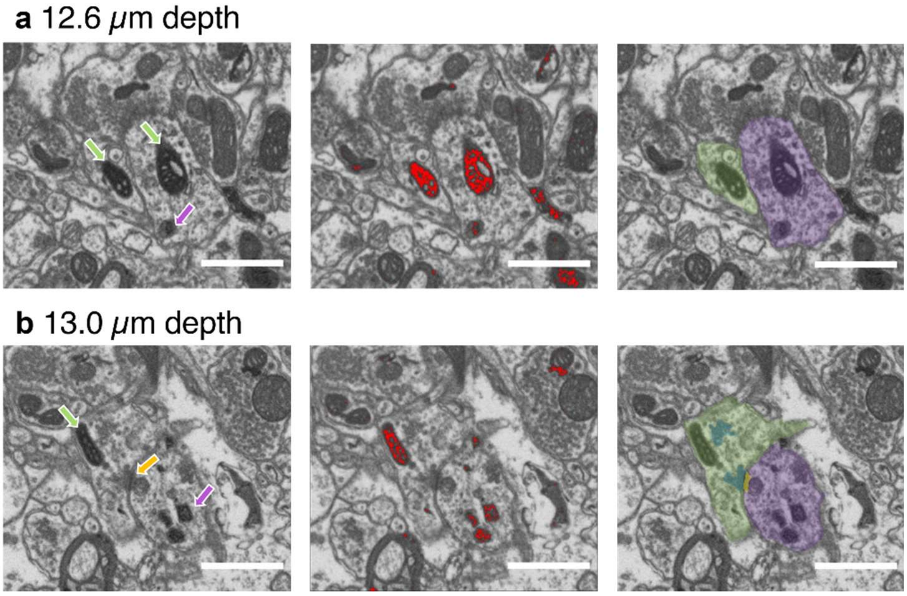


**Supplementary Figure 2 | Matrix-dAPEX2 staining is occasionally observed in dendrites. a** Left: Representative plane showing two labeled mitochondria (Sample 3). Middle: Same mitochondria are highlighted by binary mask. Right: Segmented process (green: axon; purple: dendrite). **b** Deeper plane of the same ROI showing synapse between Matrix-dApex2 labeled axon and Matrix-dApex2 labeled dendrite which also contains ER-dAPEX2 label (scale bar is 1 μm; green arrow: labeled mitochondrion; purple arrow: labeled ER yellow arrow: PSD).

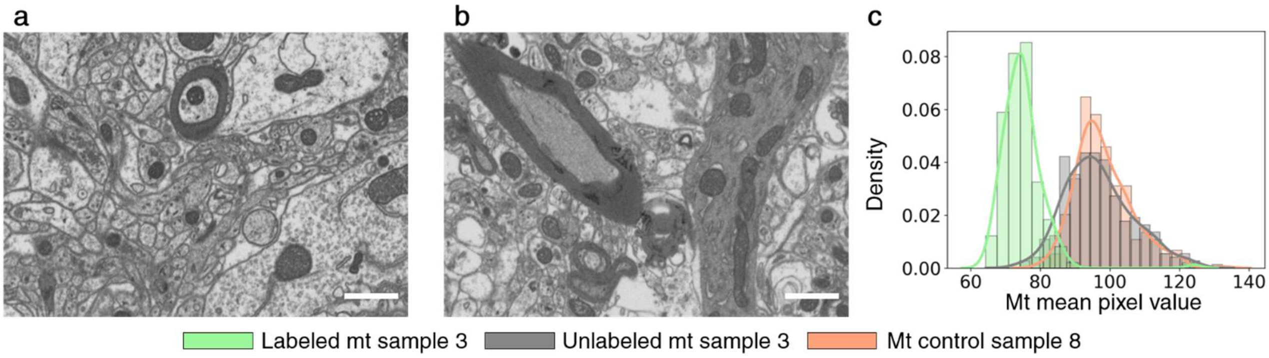


**Supplementary Figure 3 | No Matrix or ER-dAPEX2 label found in control samples.** Representative plane of control (Sample 7) showing satisfactory sample preparation quality and no dApex2 labeling. **b** Representative plane of control (Sample 8) showing satisfactory sample preparation quality and no dApex2 labeling. **c** Kernel density estimation and histogram of mitochondria mean pixel value selected from representatives XY and XZ planes in control (Sample 8) plotted with labeled and unlabeled mithocondria pixel values from a labeled sample (Sample 3; see **Fig. 1k**; scale bar is 1 μm).

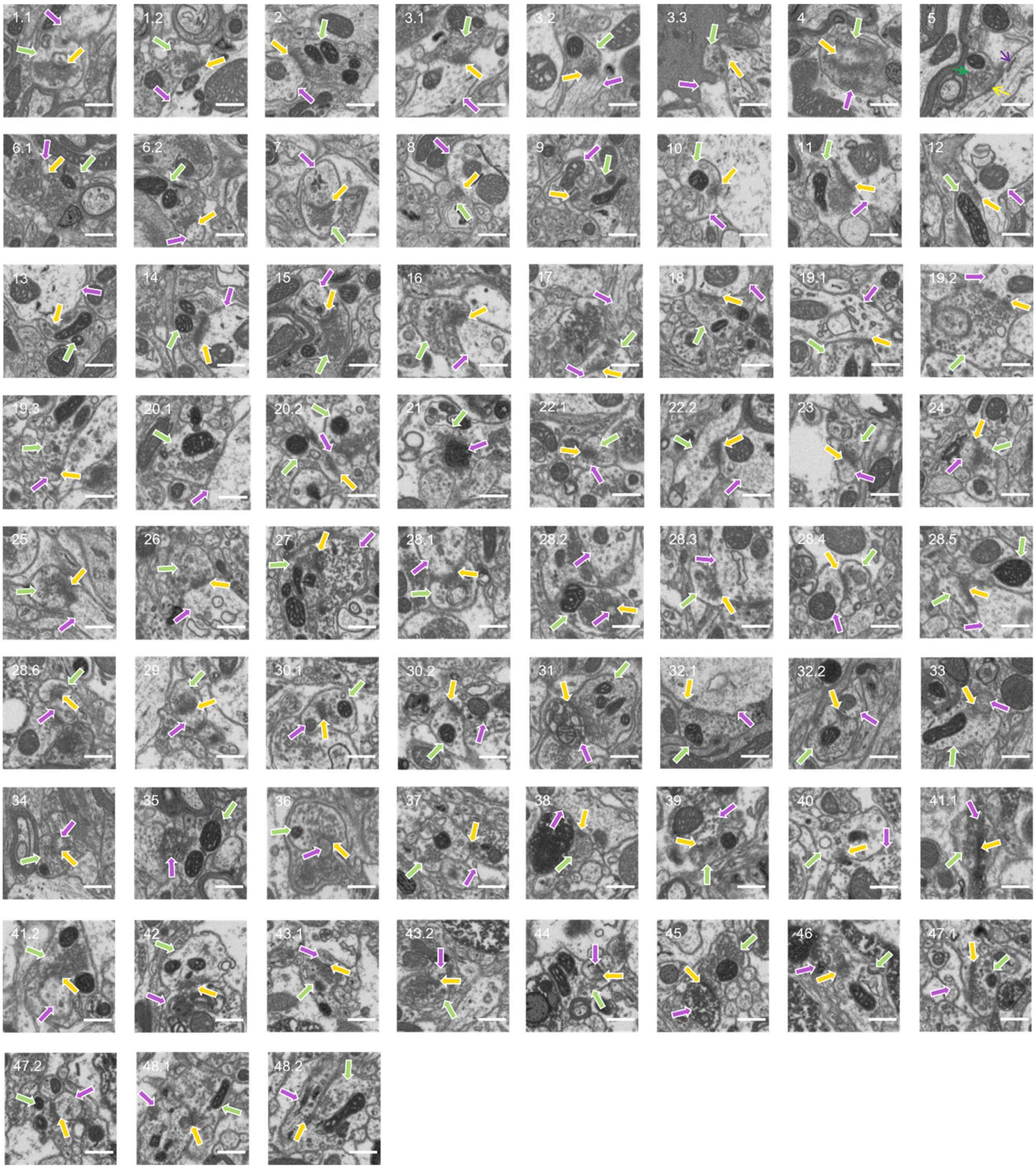


**Supplementary Figure 4 | Library of sixty-seven analyzed ACC contact sites.** (scale bar is 1 μm; green arrow: presynaptic cell with labeled mitochondrion; purple arrow: postsynaptic cell; yellow arrow: PSD).

**Supplementary Table 1 | List of samples**

| Sample ID | Animal ID | Animal genotype | Label configuration | Microscope acquisition | XYZ resolution |
|---|---|---|---|---|---|
| Sample 1 | 5460 | *Vglut2*::Cre | ACC: Matrix-dAPEX2 PAG: DIO-ER-APEX2 | TEM | 5x5 nm |
| Sample 2 | | | | SBEM | 10x10x40 nm |
| Sample 3 | | | | FIBSEM | 10x10x10 nm |
| Sample 4 | 5603 | | | | |
| Sample 5 | 7390 | | | | |
| Sample 6 | 7930 | *Vgat*::Cre | | | |
| Sample 7 | OBO-017266 | *wild-type* | Control – no surgery | | |
| Sample 8 | 8845 | *Vglut2*::Cre | | | |

**Supplementary Table 2 | Average number of mitochondria per axon in FIBSEM samples**

| Sample ID | Mean | SD | Number of axons |
|---|---|---|---|
| Sample 3 | 4.2 | 2.7 | 15 |
| Sample 4 | 2.3 | 1.5 | 6 |
| Sample 5 | 2.3 | 1.2 | 12 |
| Sample 6 | 3.8 | 3.6 | 15 |
| Total | 3.3 | 2.7 | 48 |

**Supplementary Table 3 | Summary of *Vglut2*+ dendrites and somas reconstructed in sample 3**

| Dendrite # | # Dendritic spines | # ACC contacts |
|---|---|---|
| 1 | 1 | 2 |
| 2 | 3 | 1 |
| 3 | 1 | 0 |
| 4 | 0 | 0 |
| 5* | 0 | 0 |
| 6* | 0 | 1 |
| 7* | 0 | 0 |
| 8 | 0 | 0 |
| 9 | 0 | 0 |
| 10 | 0 | 0 |
| 11 | 2 | 0 |
| 12* | 0 | 0 |
| 13* | 0 | 0 |
| 14 | 0 | 0 |
| 15 | 2 | 2 |
| 16 | 0 | 0 |
| 17 | 0 | 0 |
| 18 | 1 | 3 |
| 19 | 0 | 0 |
| 20 | 0 | 1 |
| 21 | 0 | 0 |
| 22 | 0 | 0 |
| 23 | 0 | 0 |
| 24* | 0 | 0 |

*Neuron soma was present in the block
